# Evolutionary insights from profiling LINE-1 activity at allelic resolution in a single human genome

**DOI:** 10.1101/2022.11.17.516944

**Authors:** Lei Yang, Genevieve A. Metzger, Ricky Padilla Del Valle, Diego Delgadillo Rubalcaba, Richard N. McLaughlin

## Abstract

Transposable elements including LINE-1 (Long INterspersed Element-1) impact genome variation, function, regulation, and disease. LINE-1s seem to have expanded as distinct consecutive lineages, but the drivers of lineage emergence and disappearance are unknown. Reference genomes provide a snapshot of LINE-1 evolution; however, the ongoing retrotransposition of LINE-1s in humans is not evident in these mosaic assemblies. Utilizing long-read-based haploid assemblies, we identified the sequence and location of all the youngest LINE-1s in these genomes at allelic resolution. We cloned and assayed the *in vitro* retrotransposition activity of the subset of LINE-1s with intact open reading frames and found 34 were measurably active. Yet, among individuals, these same LINE-1s varied in their presence, allelic sequences, and activity. Using a measure of *in vivo* retrotransposition of closely related groups of LINE-1, we found that recently retrotransposed LINE-1s tend to be active *in vitro* and polymorphic in the population. However, for a considerable number of LINE-1s, their measured *in vitro* activity and inferred *in vivo* fitness were uncorrelated, regardless of their frequency in the population. Some of these unexpected patterns come from rare allelic forms of old LINE-1s that retain activity, suggesting older LINE-1 lineages can persist much longer than expected. Finally, some LINE-1s showed mutations that were potentially adaptive, increasing their replication in the genome. These key mutations specific to LINE-1s with both *in vitro* activity and *in vivo* fitness represent promising candidates for the future mechanistic investigation of the drivers of LINE-1 evolution which may contribute to disease susceptibility.

## Introduction

Genomes are plagued by both infectious and endogenous parasites. Among these interlopers, transposable elements have left an indelible mark on the human and most other genomes by their recurrent and persistent integration-coupled replication [1,2]. For example, transposable elements created more than half of the modern human genome and more than 90% of some other vertebrate genomes [3]. The Long INterspersed Element-1 (LINE-1 or L1) retrotransposons are the most prolific contributors of new sequences in the human genome. LINE-1s are the only group of autonomous elements with ongoing detectable activity in human genomes [4]. As a result of their propensity for insertional mutagenesis, creation of structural variation, and ability to regulate genes, LINE-1s contribute to a slew of human diseases [5,6].

To maintain their fitness, LINE-1s must make enough new copies in the germline genome to outpace their rate of acquiring inactivating mutations. Unlike infectious retroviruses, which transmit horizontally and integrate primarily in somatic tissues, LINE-1s (and other endogenous retroelements) only transmit vertically and must, therefore, retrotranspose in the germline. This obligatory germline integration leaves a genomic fossil record of LINE-1 retrotranspositions in the genome of each host and their descendants. As a result, hundreds of thousands of LINE-1 remnants make up ~17% of human genomes and encapsulate the recent and ancient evolutionary history of these elements [1,2,4,7]. For example, LINE-1s that were active in the last mammalian common ancestor over 100 million years ago are still clearly identifiable in extant human genomes [8]. Phylogenetic analyses of these and other ancient sequences suggest that LINE-1 evolution in the ancestors of humans has been typified by a cycle of emergence, expansion, and death of LINE-1 lineages, with the downfall of an old lineage following the emergence of a new lineage [8]. One model suggests that competition among contemporaneous LINE-1 subfamilies could underlie these waves [9], but the drivers of these evolutionary transitions remain largely unknown.

In addition to sequences created by ancient (now dead) LINE-1s, the human genome also contains young and currently active LINE-1s [10–12]. These sequences reflect the most recent bouts of activity and evolution within human LINE-1s. Previous analyses estimated that among the approximately one million identified LINE-1 sequences in the reference human genome, only 3,000-5,000 are full-length (> 6 kb) [4,13]. Subsequent studies found that a large portion of the structural variation among human genomes traces back to the of activity LINE-1s including many polymorphic mobile element insertions (MEIs) within the human population [14–19]. Using both whole genome and more targeted sequencing approaches [20–24], these studies focused largely on variation in the presence/absence state (also known as ‘insertion polymorphisms’) of MEIs including LINE-1s in individual genomes. Some groups have reported hemizygosity or even sequence differences between the alleles of a specific LINE-1 insertion [25–27]. However, accurate sequencing of polymorphic LINE-1s within individual genomes has proven difficult, largely due to the fact that reads shorter than the length of LINE-1 cannot be confidently assigned to a specific locus or allele [27].

Intriguingly, some recent datasets have successfully used high-coverage long-read sequencing to determine the polymorphic state and sequence of all LINE-1s in the genome of a homozygous cell line derived from a complete hydatidiform mole (CHM) [18]. These represent the first scalable methods to catalog LINE-1 locations and sequences in individual human genomes.

Even with an accurate list of all LINE-1 sequences in an individual genome, a LINE-1’s sequence alone is currently insufficient to predict its *in vitro* or *in vivo* retrotransposition activity. Previous studies used an early draft of the human reference genome as a guide for identification of full-length LINE-1s from the assembly which were subsequently cloned and tested for their ability to retrotranspose *in vitro*. They found that only six of the cloned LINE-1s were highly active or ‘hot’ *in vitro*, suggesting a small set of young (usually polymorphic) LINE-1s generate the vast majority of the retrotransposition activity in this genome [11]. The subsequent identification of numerous population- or individual-specific, highly active LINE-1s [11,20] suggested this estimate of six ‘hot’ LINE-1s per genome may not reflect the diversity of LINE-1s and their activity in more diverse populations and individuals. However, these early studies mostly focus on the activity of a small number of LINE-1s in a small number of individuals. However, a later study identified sequence variation in the same LINE-1 across genomes that profoundly affected *in vitro* retrotransposition activity [26]. It follows that even knowing all the LINE-1 locations and sequences is not sufficient to determine the identity and activity load of all hot LINE-1s in a person’s genome.

One lofty goal of the human genetics field is to predict the ‘risk score’, reflecting the activity of LINE-1s found in an individual genome. In some cases [28–30], specific LINE-1 sequences at specific genomic locations have been shown to be causal for certain cases of human disease. The highly polymorphic state of LINE-1s among humans suggests the presence of specific disease-causing LINE-1s also varies among individuals. A fully resolved individual genome sequence could be used to identify the specific locations and sequences of LINE-1s that might predispose an individual to specific diseases. To achieve such a precise quantification of LINE-1 activity and its predicted risk would require two major advancements – first, the ability to catalog the precise location and sequence of each LINE-1 in an individual’s genome, and second, the ability to predict the activity of LINE-1s based on their sequence and location.

In this paper, we have defined the complete repertoire of intact and retrotransposition-competent LINE-1s in a single human genome. Our approach identified a higher number of active LINE-1s than previous approaches with related genomes. We used existing databases of structural variation to show that many highly active LINE-1s are polymorphic in the human population, but some are fixed, suggesting they have persisted in their active state since the last common ancestor of humans. We further demonstrate that some groups of young and old LINE-1s that are active *in vitro* have also recently retrotransposed in humans, supporting their activity *in vivo*, as well. In some cases, active LINE-1s vary in their sequence and activity among humans, suggesting that each individual may have a divergent set of highly active old LINE-1s in their genome which varies greatly among individuals. Finally, we identified sequence changes that correlate with the *in vivo* activity of certain groups of LINE-1s, some of which may represent determinants of persistent activity or targets of host restriction mechanisms. These findings demonstrate that LINE-1 polymorphisms (hemizygosity and allelic variation) are more complicated than previously thought, and hence the reference genome does not encompass the complete set of active LINE-1s in any individual. Defining the set of active LINE-1s in single haploid genomes will allow us to further define the selective pressures and evolutionary trajectories of LINE-1s in human genomes, and to lay the foundation for predictive power of activity based on sequence data.

## Results

### Comprehensive catalogue of intact LINE-1s in homozygous human genomes

The CHM1 (complete hydatidiform mole 1) assembly [31] represents the nearly homozygous genome (<0.75% heterozygosity) [32] of a human hydatidiform mole cell line derived from a European individual. Complete hydatidiform moles form from an ovum that contains no maternal DNA which is fertilized by a sperm. A single replication yields a genome homozygous for the paternal genotype. The genome of this human cell line was deeply sequenced with PacBio reads (54x), and the location and type of structural variants, including LINE-1s, found in the resulting assembly have been extensively annotated [31–33]. These published analyses mostly focused on the presence of LINE-1 insertions relative to the GRCh38 human reference genome, but the nature of these cell lines and assemblies enabled us to confidently retrieve the sequence of each LINE-1, which was not necessarily possible with previous assemblies. To collect the complete set of all LINE-1 sequences in CHM1, we extracted all sequences marked as LINE-1 from the RepeatMasker [1] annotations of the assembly. We found 919,967 sequences annotated as LINE-1, comprising ~504 Mbp of total LINE-1 sequence. This corresponds to ~16.82% of the total genome CHM1 assembly, comparable to previous estimates of LINE-1-derived sequence in GRCh38 (~one million LINE-1s and ~540 Mbp) [4,27].

Active LINE-1 sequences contain two open reading frames (ORFs) which encode proteins (ORF1p and ORF2p) required for replication (16). We assume that full-length and intact ORFs are necessary but not sufficient for LINE-1s to be active. To identify putatively-active LINE-1s in the CHM1 genome, we determined the longest continuous ORFs present in each LINE-1 sequence greater than 5,000 bp that aligned to sequences of ORF1p and ORF2p from a reference LINE-1, L1_RP_ (GenBank accession number AF148856) [29] (Figure S1 and S2). Most LINE-1s longer than 6,000 bp encode truncated ORFs, but many retain ORFs of the same length as their reference ORFs in L1_RP_ (338 codons for ORF1 and 1,275 codons for ORF2; Figure S2C). However, we also included sequences with different ORF lengths that still align along the entire length of the amino acid sequence of the reference, from start to stop codon without terminal deletions or extensions. In this way, we defined ‘intact’ LINE-1s as sequences greater than 5,000 bp which contain two intact open reading frames that align, when translated, to the full length of L1_RP_ ORF1p and ORF2p. Using this definition, we provisionally identified 148 intact LINE-1s in the assembly of the CHM1 genome.

A second human hydatidiform mole cell line (constructed similarly to CHM1) has been sequenced with 52x PacBio read depth, and the resulting assembly (CHM13) has also been extensively analyzed for structural variants [31,35]. This mole is from an individual of unknown ethnic origin, but clusters with CHM1 and other European genomes. We applied our computational pipeline to identify intact LINE-1s in CHM13. The distribution of LINE-1 sequence lengths in CHM13 was similar to that of CHM1, and we identified 142 intact LINE-1s in the CHM13 assembly (Figure S3).

Sequencing and assembly errors could contribute to an underestimation of intact LINE-1s using our assumptions/pipeline. In particular, PacBio sequencing errors most often occur as indels in homopolymer tracts [36], a sequence pattern present at several locations in the LINE-1 sequence, which could results in discarding of bona fide intact sequences. To assess the prevalence of sequencing errors that disrupted true intact LINE-1s, we identified sequences that deviated from an intact LINE-1 sequence by a single frame-shifting mutation (82 LINE-1s) or two frame-shifting mutations (47 LINE-1s). We assumed that frame-shifting mutations shared by any two assemblies (CHM1, CHM13, hg38) were not likely to be the result of sequencing errors and identified 27 of these LINE-1s with frame-shifting mutations in that were unique to CHM1 (24 with single frame-shifts and 3 with two frame-shifts). We took advantage of a publicly available BAC library of the CHM1 genome (CHORI-17, The BAC clones from the hydatidiform mole were created at BACPAC Resources by Drs. Mikhail Nefedov and Pieter J. de Jong using a cell line created by Dr. Urvashi Surti. available at BPRC, https://bacpacresources.org) to Sanger sequence these apparently frame-shifted LINE-1s. While most of the frame-shifting mutation present in the assembly were confirmed, we identified sequencing errors in four CHM1 LINE-1s with one frame-shifting mutation in the published CHM1 assembly which revealed them to be intact LINE-1s. None of the LINE-1s containing two annotated frame-shifting mutations in the assembly were intact by Sanger sequencing, so we did not pursue further sequencing of LINE-1s containing two or more frame-shifting mutations. In a complementary analysis of the CHM13 genome, we leveraged a new, high fidelity CHM13 assembly based on circular consensus long reads [35]. Using this high-fidelity assembly to check the sequence of 87 LINE-1s in the original CHM13 assembly containing one frame-shifting mutation and 45 LINE-1s containing two frame-shifting mutations, we identified two additional intact LINE1s (both from the single-shift mutation pool).

In addition to frame-shifting sequencing mutations, we also expected evidence of assembly errors or gaps that excluded intact LINE-1s from the CHM assemblies. Since the donors for CHM1 and CHM13 genomes clustered together with other European genome, we expected LINE-1s present in only one of these two, closely-related genomes to have low or intermediate frequencies in the human population. However, we observed two LINE-1s that were present in the CHM13 assembly, absent from the CHM1 assembly, and apparently present in all other human assemblies in the KGP [37]. Indeed, these LINE-1s fell in regions of the assembly with a high density of contig junctions; Sanger sequencing of these regions from corresponding BACs found that contrary to initial observations relying on assembly sequence, both of these LINE-1s are indeed present and intact in CHM1. The flanking sequence of some intact LINE-1s could not be located in the reference genome (GRCh38) using the UCSC genome browser liftover tool. We were also unable to find these unplaced LINE-1s a more complete assembly of this genome – the draft telomere-to-telomere (T2T) CHM13 assembly [38–40]. Together, our detailed re-analysis of intact LINE-1s in multiple assemblies and targeted resequencing of LINE-1s identified 154 and 144 intact LINE-1s in CHM1 (File S1) and CHM13 (File S2), respectively, by cross-referencing and resequencing suspected errors and gaps.

### A pseudo-diploid genome reveals LINE-1 allelic variation

LINE-1 insertions, especially young insertions, are highly polymorphic in the human population [22,41–43]. Some LINE-1 insertions are also hemizygous, meaning that LINE-1 is only present at a given locus in one of the two chromosomes/haplotypes in an individual. In individuals that are homozygous for a given LINE-1 insertion, variation in the sequence of the two alleles of that LINE-1 has also been reported [25,26]. Clearly LINE-1s are a major contributor to human genome variation; however, we lack a thorough understanding of the scale of LINE-1 insertion and allelic heterozygosity within and between individual genomes. We used the CHM1 and CHM13 assemblies to make genome-wide estimates of the content and variation of LINE-1 that could exist within a single diploid genome. In total, we found a combined 294 intact LINE-1 ‘alleles’ present at 194 distinct loci in the union of these two haploid genomes (Table S1). Most of the intact LINE-1s were present (sharing synteny but not necessarily intactness) in both CHM genomes (131/154 in CHM1 and 121/144 in CHM13; Figure 1A). Of these shared LINE-1s, 102 were intact in both CHM1 and CHM13 (Figure 1A, purple shaded area). Twenty-nine LINE-1s that are intact in CHM1 are present in CHM13 but have accumulated ORF-disrupting mutations, and 19 LINE-1s are intact in CHM13 but contain ORF-disrupting mutations in CHM1 (Figure 1A). Finally, there are unique LINE-1 insertions in each genome – 23 in CHM1 and another 23 in CHM13 (Figure 1A). Each of these loci is hemizygous (with an intact LINE-1 in one genome and devoid of a LINE-1 insertion in the other genome) and likely represent the youngest LINE-1s in these two genomes.

**Figure 1:**
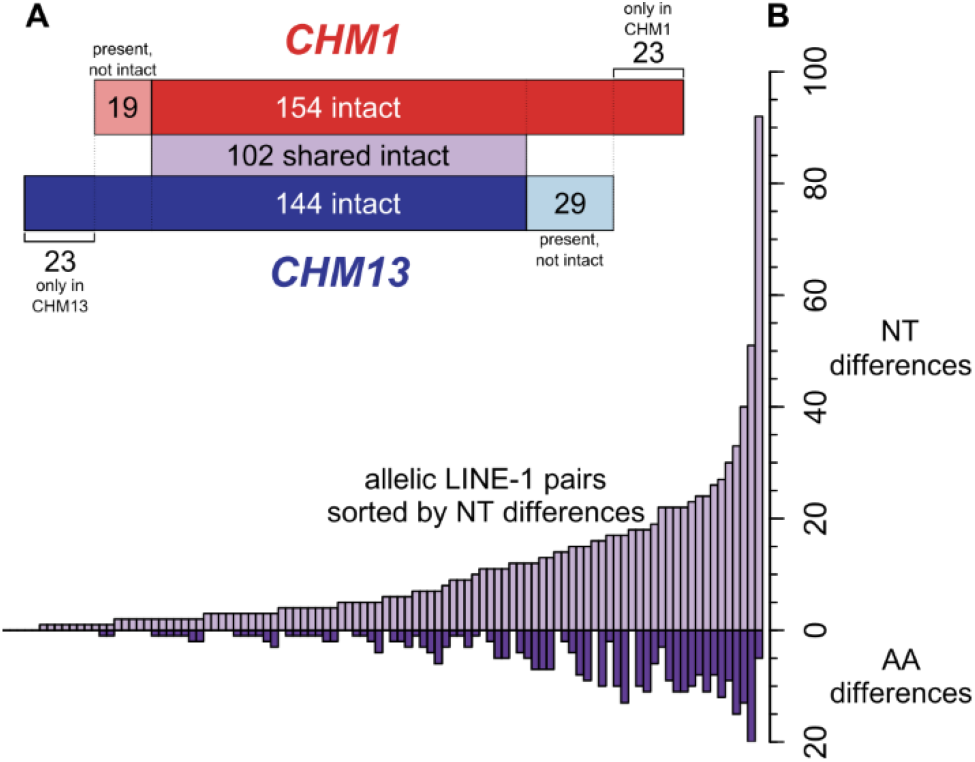
A comparison of intact allelic LINE-1 pairs across two haploid genome assemblies. A) A comparison of the intact LINE-1s in two nearly homozygous genomes (CHM1, top/red, and CHM13, bottom/blue) based on liftover found 102 shared intact LINE-1s (light purple shaded area). Of the LINE-1s that are intact in only one genome, some are present but not intact in the other genome (light red and light blue shaded areas), and some are absent in the other genome. B) Distribution of nucleotide (top) and non-synonymous (bottom) changes between CHM1 and CHM13 of the 102 allelic intact LINE-1s ordered by nucleotide differences. NT: nucleotide; AA: amino acid.

We were also able to resolve sequence differences at the 102 loci that contained intact allelic pairs of LINE-1s in both CHM1 and CHM13. Some diversity of LINE-1 alleles at a single locus has been described [25], but the CHM assemblies provided an unprecedented opportunity to learn about this form of diversity across an entire genome. At the nucleotide level, some of the largest differences within pairs arise from large deletions in the UTRs; these are scored as multiple single nucleotide differences in the alignment, but often represent single deletion events. Other pairs have deletions and SNPs distributed throughout the UTRs and coding sequence. Only 4/102 allelic pairs of LINE-1s are identical at the nucleotide level between CHM1 and CHM13 assemblies, while 63/102 pairs are identical at the amino acid level. The most distinct intact LINE-1 allelic pairs differ by 20 amino acid- and 92 nucleotide changes, respectively (Figure 1B). We confirmed the sequence of the CHM1 version of four of these highly divergent LINE-1 pairs by resequencing regions of CHM1 BACs containing these LINE-1s. We focused on the four CHM1 LINE-1s containing the greatest number of differences relative to its partner in the CHM13 assembly. In each case, our re-sequencing perfectly matched the assembly sequence we had analyzed, suggesting that these allelic differences were unlikely to result from sequencing errors. Together, our results revealed pervasive allelic differences among the LINE-1s in the two CHM genomes studied.

### Comprehensive measurement of LINE-1 in vitro activity in a human genome

One additional outstanding question in the LINE-1 field has been the true number of active LINE-1s within a single genome. Measurements based on the draft human genome reference found six highly active LINE-1s, though it is now clear that version of the reference genome was biased towards high frequency, older LINE-1s. Subsequent studies targeting LINE-1s not present in the human reference genome found many active elements among six diverse human genomes and estimated fourteen highly active LINE-1s in one of these individuals. The CHM1 genome and its associated BACs and their mapping to the GRCh37 reference genome (available at BPRC, https://bacpacresources.org) have made it possible for us to survey the number of active LINE-1s within a single genome with unprecedented resolution. As a first step, we set out to measure the *in vitro* retrotransposition activity of all intact LINE-1s in the CHM1 genome, by cloning them from BACs and testing their activity in an established cell-based assay [34,44]. We cloned 142 intact LINE-1s by PCR amplifying the complete LINE-1 sequence (including UTRs) as defined by RepeatMasker and inserting them into a retrotransposition vector with a luciferase reporter. We then transfected three independent clones of each LINE-1 into 293T cells and measured the normalized luciferase signal to determine the retrotransposition level of each clone. We compared retrotransposition of each test clone to a commonly studied and highly active human LINE-1, L1_RP_ [29]. As additional controls, we also included a mutant LINE-1 lacking activity (JM111, ORF1p R261A/R262A [34]) and an empty vector without a LINE-1. Notably, there were nine intact LINE-1s that we were unable to map to a CHM1 BAC (many due to their location within centromeric or simple repeats), and three other intact LINE-1s that we failed to successfully clone.

We observed that the majority of the intact LINE-1s from CHM1 had no detectable *in vitro* retrotransposition activity, consistent with previous activity studies [11,20] and our understanding of LINE-1 mutation accumulation. With our CHM1-based approach, we found 34 LINE-1s with measurable retrotransposition activity (>5% L1_RP_; 27 LINE-1s were >10% L1_RP_). Of these 34 elements, two had significantly higher activity than L1_RP_ and three elements had activity comparable to that of L1_RP_ (Figure 2). These data suggest that human genomes contain a large pool of highly active LINE-1s and that previous reference genomes were biased against the youngest, polymorphic LINE-1s in the population.

**Figure 2:**
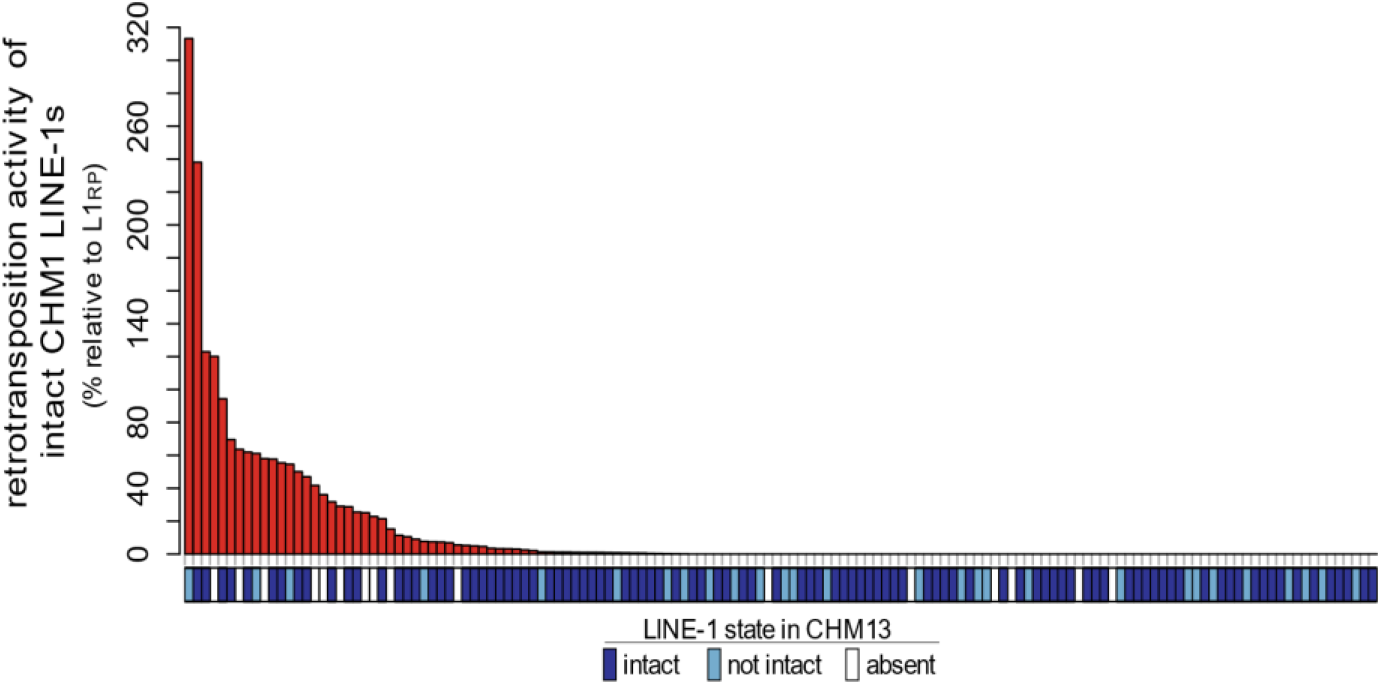
*In vitro* retrotransposition activity of all intact LINE-1s in a haploid genome. Red bars show the retrotransposition activity of each intact LINE-1 from the CHM1 genome normalized to the positive (L1_RP_: 100%) and negative control (empty vector: 0%). A LINE-1 is considered ‘active’ only when it is significantly more active than the negative control (*p* < 0.01, student’s t-test) and its activity is greater than 5% that of the positive control. Blue/white boxes below each bar shows whether each intact LINE-1 from CHM1 is present/absent and intact/not-intact at the syntenic allele in CHM13.

We then turned to a detailed comparison of the sequence of the 34 active LINE-1s from CHM1 in another human genome. We found that 20 were intact in CHM13, four were present but not intact in CHM13 and 10 were absent from CHM13 (Figure 2). This high variation in the presence of active LINE-1s between closely related genomes further supports the notion that the repertoire of active LINE-1s varies greatly among human individuals [20,26]. It remains unclear how varied the number of active LINE-1s may be across more diverse genomes [23].

Overall, we found that the CHM1 genome contains two of the six previously described ‘hot’ LINE-1s [11]; both of these LINE-1s were also highly activity in our assays (36% and 70% of L1_RP_). We also found three recently reported intact LINE-1s that are polymorphic in humans (from Chinese and Japanese genomes; [20]) in the CHM1 genome and all exhibit high activity (123%, 47%, 25% of L1_RP_). Many LINE-1s that were highly active in our analyses were reported as weakly active or dead in another study. For example, the two most active LINE-1s in our analysis (238% and 313% of L1_RP_) were both previously studied but found to be very weakly active [11].

We hypothesized that the observed difference in activity could arise from sequence variation in these allelic LINE-1s that modulates their activity. Indeed, a previous study found extensive variation in the *in vitro* activity of the alleles of a single LINE-1 from several individuals, leading them to propose a model in which mutation accumulation gradually inactivates the LINE-1 alleles in a population [26]. To test whether allelic variation in LINE-1s may modulate *in vitro* transposition activity, we collected all available allelic forms of several known ‘hot’ LINE-1s from CHM1, CHM13, the NCBI nt database (including Brouha *et al*. [11] and Beck *et al*. [20]), and the GIAB project [45] to identify candidate mutations that are shared by alleles with similar *in vitro* activity profile, and hence might underlie the *in vitro* activity discrepancy.

We compared each pair of alleles for which our retrotransposition measurement differed from a previous measurement. We scanned for non-synonymous changes present in the less active allele that are absent from all known active alleles of that locus. Using these pairs, we defined one or two sites fulfilling these criteria which we used as surrogates of their *in vitro* activity for the alleles that are not tested in the *in vitro* activity assay (File S3). In all these instances, we found that the allele with *in vitro* activity was the minor allele. This suggests that determination of the active LINE-1s from one genome is not sufficient to accurately predict the activity of even an identical set of insertions in another individual. Instead, each genome likely harbors several LINE-1s that exist in their active state in only a small fraction of individuals (the rare/minor allele). These data suggest that non-synonymous changes in intact allelic insertions drive the variation in the active set of LINE-1s even between two closely related individuals.

### Frequency spectrum of all LINE-1s in a single genome

The bulk of LINE-1 activity in human genomes is thought to originate from young, polymorphic LINE-1s [11]. Empirically, we found that 33/34 of the *in vitro* active LINE-1s in CHM1 come from the most recently expanding groups of the human-specific LINE-1s (L1Hs). Ta1d, the youngest subfamily of L1Hs, accounts for the largest fraction (15/34) of *in vitro* active elements; we also detected *in vitro* activity from each of the older subfamilies of L1Hs: Ta1nd (3/34), Ta0 (9/34), and nonTa (6/34). Unexpectedly, we found that one of the *in vitro* active LINE-1s belongs to the L1PA2 group, a hominid specific group, suggesting that this element predates the last common ancestor of humans, and challenging the previous observations that all current LINE-1 retrotransposition comes from the youngest subfamilies.

Within the L1HS elements, it is thought that the youngest, polymorphic LINE-1s are most likely to be active. Given our *in vitro* activity and sequence data for active LINE-1s, we could more rigorously look for correlations between population frequency and *in vitro* activity. To do this, we determined the ‘allele frequency’ in the human population of all intact LINE-1s found in the CHM1 and CHM13 genomes. We integrated data from the thousand genome project (KGP) [37], euL1db [46], and two previously published studies [22,47] (Table S2, column V). From this survey, 142/154 intact CHM1 LINE-1s had available data for both population frequency and *in vitro* activity. More than half of the intact CHM1 LINE-1s (74/142) are fixed in the human population; the remaining active polymorphic LINE-1s are relatively evenly split between the rare and common insertions in this genome (Figure 3, top histogram). Of interest, seven LINE-1s are only found in CHM1, but not in the related CHM13 genome, suggesting these insertions are very young.

**Figure 3:**
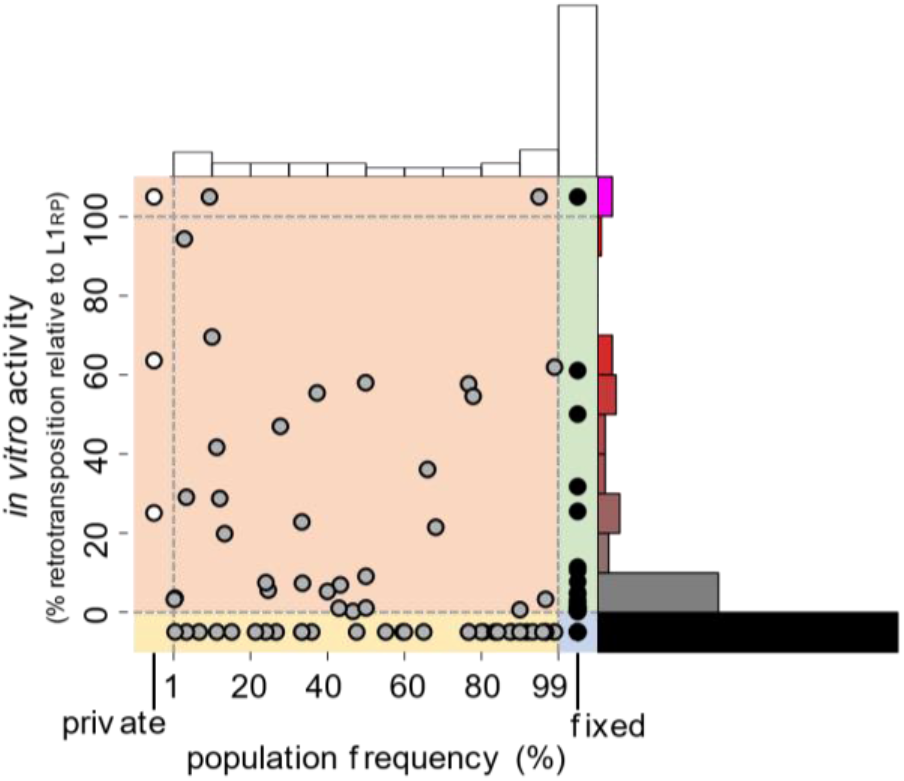
Comparison of *in vitro* activity and population frequency of intact CHM1 LINE-1s. Top and right, histograms of population frequencies and retrotransposition activity of CHM1 intact LINE-1s. The scatter plot shows a dot for each LINE-1, placed according to its activity and population frequency. LINE-1s that are fixed in the human population are contained in the ‘fixed’ bin. LINE-1s with higher activity than L1_RP_ were binned together and LINE-1 with activity values not significantly different than a negative control were binned together for display purposes. Shaded regions within the scatter plot correspond to the combinations of fixed/polymorphic, active/not active *in vitro*, and high/low in vivo fitness as shown in Figure 5A.

Next, we compared these frequency estimates to our *in vitro* activity data. We found that almost half (67/142) of all intact LINE-1s in CHM1 are fixed in humans and inactive *in vitro* (Figure 3; rightmost bar of top histogram). Overall, we find LINE-1s in every frequency bin that retain *in vitro* retrotransposition activity, and the most highly active LINE-1s are either very low frequency or fixed in humans. In fact, we found 8 fixed LINE-1s that were active *in vitro* (Figure 3, green highlight). Thus, for LINE-1s that are polymorphic and active *in vitro* (Figure 3, orange highlighting), population frequency and activity are not obviously correlated (R^2^ = 0.035), suggesting that while many of the retrotransposition-competent LINE-1s in each human genome are young and polymorphic (the prevailing model), a substantial number of old and high frequency LINE-1s also retain significant *in vitro* activity.

### Measuring the activity history of a LINE-1 in vivo

Evolutionary successful LINE-1s must retain *in vitro* activity but the ability to produce new copies in the germline genome (*in vivo* activity) imposes additional constraints. We reasoned that LINE-1s with many close relatives were either recently active *in vivo* or are closely related to a LINE-1 with recent *in vivo* activity. Therefore, the distribution of the sequence differences between a LINE-1 and each other full-length LINE-1 in the genome should reflect that LINE-1’s recent *in vivo* activity, and this measurable distance could be used as a proxy for the *in vivo* activity of each intact CHM1 LINE-1. We measured the Hamming distance for each intact LINE-1 nucleotide sequence in the CHM1 assembly to every other full-length LINE-1 sequence in the genome (including all sequences great than 6kb, not just intact LINE-1s). Each LINE-1 exhibited one of three broadly defined distributions which likely reflect undetectable, ancient, or recent *in vivo* activity (Figure 4A). The pairwise distances for a given LINE-1 form some combination of three apparent peaks, which represent its close, intermediate, and distant relatives. All LINE-1s are distantly related to some set of the other LINE-1s in the genome, and these relationships are reflected in the peak of pairwise distances greater than 82 nt substitutions that appears in every distribution (Figure 4A, ‘old’ bin). LINE-1s with a distribution containing only this ‘old’ peak had no close relatives and hence no detectable *in* vivo activity (Figure 4A, black histogram). Some LINE-1s had a distribution with a second peak spanning 28-82 nt substitutions (Figure 4A, ‘mid’ bin). This type of distribution arises from older *in vivo* activity which resulted in detectable but ancient expansion of that LINE-1 or its close relative (Figure 4A, gray histogram). Finally, some LINE-1s had three peaks – an ‘old’ peak, a ‘mid’ peak, and a third peak which comes from close relatives with only 1-27 nt substitutions (Figure 4A, ‘young’ bin). The presence of this group of highly similar LINE-1s suggest recent *in vivo* activity of these LINE-1s or their close relatives in the ancestral lineage of CHM1 (Figure 4A, red histogram). We use the number of LINE-1s found within this young, closely related region of the distribution as a proxy for the recent i*n vivo* activity of each LINE-1 (Figure S4). Thus, by measuring the activity history of all intact LINE1s in the CHM1 genome, we found that most intact LINE-1s have ‘no near neighbors’ (114/142 with black- or gray-type distributions), which supports the conclusion that the majority of the intact LINE-1s in that genome are not active *in vivo*. Yet, 28/142 LINE-1s show evidence of recent *in vivo* activity by them or a close relative, seemingly comparable in scale to the 34 LINE-1s with detectable *in vitro* activity in our retrotransposition assay. However, LINE-1s with high *in vitro* activity are not always also high *in vivo* activity (Figure 4B). We did observe many LINE-1s that showed, expectedly, low *in vitro* and *in vivo* activity, and another group with high *in vitro* and *in vivo* activity. In addition, we found many outliers to this expected correlation which suggest these LINE-1s have experienced more complex evolutionary histories (see Discussion).

**Figure 4:**
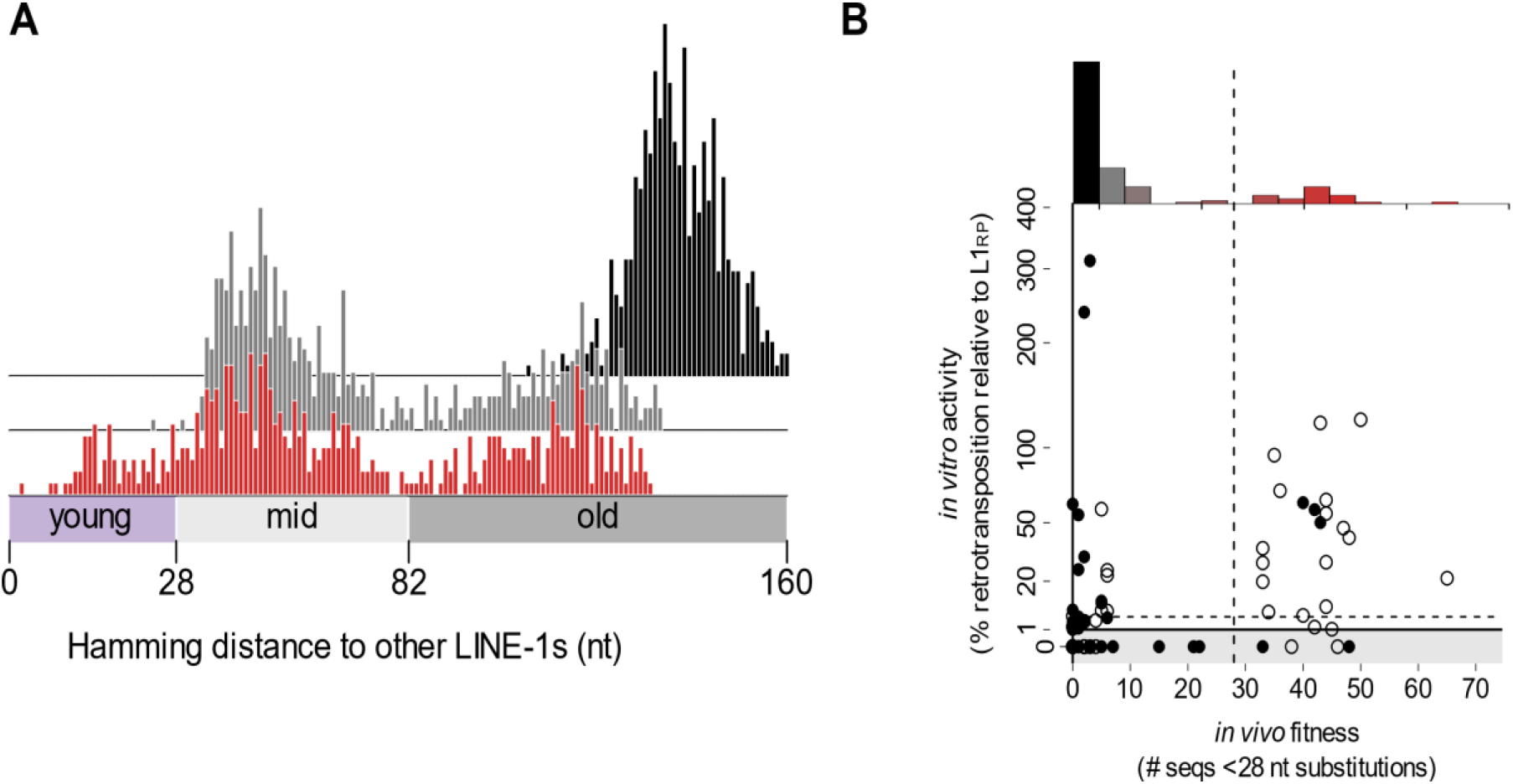
A measure of *in vivo* LINE-1 fitness. A) Examples of the distributions of sequence distance of individual LINE-1s to all other full-length LINE-1s in CHM1, used to infer the *in vivo* fitness of a LINE-1 and its close relatives. A young (red), a mid-age (gray), and an old (black) example LINE-1 are shown. B) Scatter plot of *in vitro* activity versus *in vivo* fitness of each intact CHM1 LINE-1. Top, histogram of *in vivo* activities of intact LINE-1s in CHM1. Filled dots represent the frequent (> 75%) LINE-1s in the human population, and hollow dots represent the polymorphic (≤ 75%) LINE-1s in the human population. LINE-1s with *in vitro* activity smaller than or equal to negative control are separately plotted in the gray area of the plot. Dashed lines represent the cutoffs used to call *in vitro* active/inactive and *in vivo* fit/unfit.

### An analysis of LINE-1 evolutionary history by integration of population frequency, in vitro activity, and in vivo activity

Although we predicted that the pairwise comparisons of population frequency versus *in vitro* activity and *in vitro* versus *in vivo* activity would show correlation for most LINE-1s, our data show that this is not necessarily true for certain LINE-1s. We predicted that a 3-way comparison of these parameters might explain some outliers within each pairwise comparison. Based on the distribution of population frequency (Figure 3, top), *in vitro* activity (Figure 3, right), and *in vivo* activity (Figure 4B), we binarized each distribution into polymorphic (<= 75% frequency in the human population), frequent (> 75% frequency) *in vitro* active (>5% *in vitro* activity of the positive control), *in vitro* inactive (<=5%), *in vivo* fit (<=10 near neighbors) and *in vivo* unfit (>10 near neighbors) categories. Based on the combination of these parameters, the intact CHM1 LINE-1s were spread across all eight possible categories (Figure 5A).

**Figure 5:**
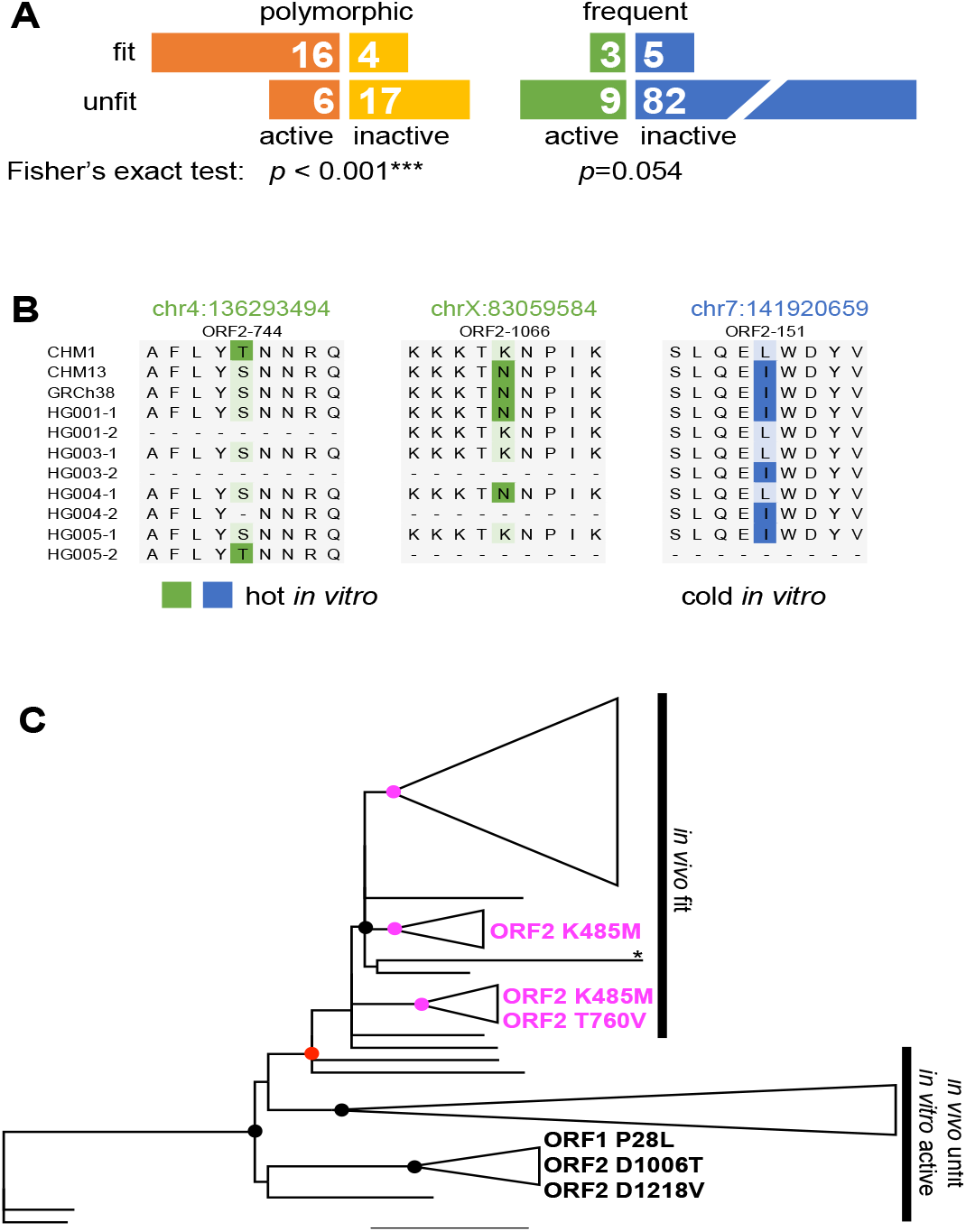
Discrepancy between *in vivo* and *in vitro* activity suggests recent adaptation of LINE-1s. A) Contingency table showing the number of LINE-1s in each category: polymorphic: ≤ 75 % frequency in the human population; frequent: > 75% frequency in the human population; active: > 5% *in vitro* activity of L1_RP_; inactive: ≤ 5% *in vitro* activity of L1_RP_; fit (*in vivo*): >10 near neighbors (≤ 28 substitutions); unfit (*in vivo*): ≤ 10 near neighbors (≤ 28 substitutions). B) Candidate amino acid changes responsible for *in vitro* activity discrepancy between different allelic forms at one example site. Coordinates indicate the start of the corresponding LINE-1 in the GRCh38 reference genome. ‘HG00X’ indicates the sample number from the GIAB project, ‘-1’ and ‘-2’ indicate the two alleles from the same diploid genome. Continuous ‘-’s indicates that LINE-1 is absent from the allele. C) Maximum likelihood phylogeny of the *in vivo* unfit LINE-1s and the *in vivo* fit, *in vitro* active LINE-1s. Dots on the backbone mark the supported nodes (aLRT ≥ 0.85), and the red dot indicates the node that separates the two major groups. Magenta dots indicate the supported clusters within the *in vivo* fit group. ‘*’ indicates the outlier active-unfit sequence that fell into the *in vivo* fit group. Labelled non-synonymous substitutions and their ORF coordinates that are specific to the cluster.

Predictably, more than half of all intact CHM1 LINE-1s that we successfully cloned were old and dead (82/142). These L1s were fixed or of high frequency, with no detectable *in vitro* activity, and no closely related LINE-1s in the genome (low *in vivo* activity). Our pairwise comparison showed that *in vitro* activity cannot be reliably predicted by *in vivo* activity or frequency alone. However, we found that within the set of polymorphic LINE-1s, *in vivo* and *in vitro* activity are correlated (Fisher’s exact test, *p*=6.68×10^-4^, Figure 5A, left), but not within the set of frequent or fixed LINE-1s (Fisher’s exact test, *p*=5.44×10^-2^, Figure 5A, right). These data suggested that the population frequency and *in vivo* activity could be integrated into a more accurate prediction of the *in vitro* activity of specific LINE-1s or LINE-1 alleles. In this way, the wet lab measurement of retrotransposition activity (the current benchmark for LINE-1 activity) could be predicted from two computable parameters.

Even with this statistically significant correlation, we observed several outliers with high *in vivo* fitness but no detectable *in vitro* activity. Indeed, *in vivo* fitness derives from the replication history of the collection of LINE-1 alleles at that locus in many genomes, while the retrotransposition assay records the *in vitro* activity of just one of the alleles at the locus. We posited that this discrepancy between *in vivo* and *in vitro* activity could result from differences at the amino acid level in the CHM1 allele relative to other (active) alleles in the population. To test this, we selected our *in vivo* fit and *in vitro* inactive LINE-1s that had been previously tested for *in vitro* retrotransposition by other groups [11,20]. For this set of LINE-1s, we collected all available alleles from the reference genome, GenBank and the GIAB projects [45]. Utilizing our identified surrogate sites of *in vitro* activity discrepancies (File S3 and associated text above), we showcased three LINE-1s with dramatic allelic variation of their *in vitro* activity (Figure 5B). The three examples show that both *in vitro* active and inactive LINE-1 alleles are common in the population (consistent with a slow decay of LINE-1 sequences after their insertion) [26]. Observation of these ‘hidden’ active LINE-1 alleles suggests that the activity of some apparently-dead LINE-1s is likely more persistent than we thought before because each person has a different set of active LINE-1 alleles.

Previously, *in vitro* retrotransposition activity has been used as the primary indicator of activity in humans because of the difficulty of making true *in vivo* activity measurements [48]. Our *in vivo* fitness metric required persistent *in vivo* activity of the LINE-1 or related lineage members over recent human evolution. In further support of the disconnect between *in vitro* activity and *in vivo* fitness, we found 15 retrotransposition-competent (*in vitro* active) LINE-1s with no evidence for recent *in vivo* activity (Figure 5A, the ‘active’ and ‘unfit’ LINE-1s). This could arise, for example, if a LINE-1 retains the sequence determinants of activity but inserted into a non-permissive genomic location. Alternatively, these LINE-1s could be more potently restricted by the host in the germline compared to other LINE-1s. When we generated a phylogeny to compare the ‘fit’ LINE-1s and the ‘active-unfit’ LINE-1s (Figure 5C, File S4), we found a single supported node (Figure 5C, red dot, aLRT=0.92) separated all but one (Figure 5C, marked by ‘*’) active-unfit LINE-1 from the ‘fit’ LINE-1s. This suggests specific amino acid sequences are shared within the *in vivo* fit group, distinct from the amino acids shared by the *in vivo* unfit group. Hypothetically, these phylogenetically informative sites could represent determinants of susceptibility or resistance to host restriction factors. Within the *in vivo* ‘fit’ CHM1 LINE-1s, we observed 28 fit L1s forming three distinct supported clusters (Figure 5C, magenta dots, aLRT > 0.85). Although most *in vitro* active LINE-1s belong to the youngest subfamily of the human-specific LINE-1s, Ta1d [11,20,49], these clusters suggested further diversification within the Ta1d subfamily with yet unknown consequences for potential emergence of new active LINE-1 subgroups.

## Discussion

Decades of research have bridged the first report of an active LINE-1 in humans to extensive databases of LINE-1 insertions in hundreds of human genomes [11,15,20,22,46]. Many of these datasets address the population genetics of LINE-1s in humans, but many have also analyzed the sequence and *in vitro* activity (retrotransposition rate) of select LINE-1 mobile element insertions (MEIs) from various genomes [11,20]. Our data bridges a notable gap in our current understanding of LINE-1 diversity, activity, and evolution in humans: a complete picture of the frequency, allelic diversity, and activity of the intact LINE-1s in a homozygous human genome. Because LINE-1s are the only active autonomous transposable elements in the human genome, and they are known to be causal or central to a slew of human diseases, these comprehensive analyses of all active LINE-1s in a single genome hold value for understanding and predicting with precision, the ‘genomic load’ of active LINE-1s in each individual.

Previous work towards this goal used the LINE-1s sequences represented in the most recent, but incomplete reference genome assemblies. These assemblies were also biased by the averaging effect of collapsing multiple genome sequences into a single haploid reference sequence. In this case, the non-representative genotype of donors neither faithfully represents the complexity found in the human population, nor fully describes the genome of any one individual. Our work addresses several fundamental questions regarding LINE-1 variation and recent adaptation by utilizing high quality haploid and long-read-based genome assembly. Each LINE-1 in our collection has been previously identified as a structural variant in the CHM genomes [18]. However, the curation of these sequences using a layered approach of RepeatMasker LINE-1 calls followed by in-depth analysis of sequence characteristics and their functional properties is a major step towards understanding the functional consequences of the substantial sequence variation in the LINE-1s of individuals.

By retrieving and comparing LINE-1 alleles, we show that LINE-1 alleles are variable in their sequence, ORF intactness, and *in vitro* retrotransposition activity. By measuring the *in vitro* activity of all 154 intact LINE-1s in a single haploid genome, we show that human genomes harbor more active LINE-1s than previous estimations. Motivated by previous approaches [50], we developed a simple distance-based metric to predict the historical *in vivo* retrotransposition activity of a LINE-1 and its close relatives. We found that, in general, this ‘*in vivo* fitness’ parameter only correlates with *in vitro* activity for LINE-1s that are polymorphic in the human population. The few outliers with conflicting *in vitro* activity and *in vivo* fitness demonstrate the surprising persistence of older LINE-1 lineages and adaptation of the youngest LINE-1s. These findings are foundational to our understanding of LINE-1s because evolutionary dynamics of the youngest LINE-1s are previously mostly inferred from a few fixed or high-frequency LINE-1s in the human population, and only inferred from certain allelic forms of these. Since LINE-1s are a major source of human genomic variation [14] and the cause of numerous diseases [6], such advancement in our understanding of LINE-1s is a new start for us to understand consequences of LINE-1 retrotransposition in the context of human health.

Our data reveal LINE-1 variation at multiple levels: LINE-1s at the same allele can vary by their ORF intactness, sequence, and *in vitro* activity. Previously, studies about LINE-1 variation mostly focused on their polymorphism (presence/absence) in the human population due to the limitations of using short reads to resolve the sequence and alleles of LINE-1s. The recent availability of long-read-based assemblies of the haploid CHM genomes enabled our investigation of genomic LINE-1 at allelic resolution. Previously, Lutz *et al*. [25] was the first attempt to tackle the allelic variation of LINE-1s, finding that common LINE-1 alleles at a single locus exhibited up to 16-fold differences in retrotransposition assays. Later, Seleme *et al*. [26], assaying activity LINE-1 alleles at three loci, suggested that the repertoire of LINE-1 alleles in any individual can differ dramatically in their cumulative retrotransposition rate. Our data expand upon these previous studies to include all intact and retrotransposition-active LINE-1s in a pseudodiploid genome and suggest that approximately one-third of the intact LINE-1s in one CHM genome are present but not intact in the other (Figure 1A). Further, we found pervasive variations between the two CHM genomes, including up to 33 dispersed nucleotide changes and deletions ranging from 11 bp to up to 80 bp.

Our *in vitro* retrotransposition assays showed that 34 CHM1 LINE-1s have activity greater than 5% of the activity of L1_RP_ (27 were above 10% and 24 were above 20%), compared to the previous estimates of 14 LINE-1s greater than 5% L1_RP_ activity by Brouha *et al*. [11] based on HGWD (they reported six ‘hot’ LINE-1s with above 20% L1_RP_ activity, and 11 with above 10% L1_RP_ activity). Since the CHM1 genome is not complicated by LINE-1 hemizygosity or heterozygosity, one could assume a simple additive model to estimate that 68 LINE-1s with measurable *in vitro* activity reside in this diploid human genome. This number is comparable to the inference of 80-100 active LINE-1s per diploid genome made by Brouha *et al*. [11]. We expect this number to vary substantially amongst individuals based upon the number and age of LINE-1s in the underlying populations.

This empirical assessment of active LINE-1s in a human genome being closer to previous inferences likely derives from our analysis of the more complete haploid-based assembly. Although fewer active LINE-1s were found in HGWD, it is not simply a subset of the active LINE-1s found in CHM1: CHM1 and HGWD only share two LINE-1s with higher than 5% activity of L1_RP_. Further comparison between our two haploid genomes also confirms such variation: 14 of the 34 active CHM1 LINE-1s are either not intact or not present in CHM13 (Figure 2). Based on the comparison between the two haploid genomes and HGWD, we expect the set of *in vitro* activity LINE-1s to vary substantially amongst individuals. Indeed, the investigation of the LINE-1 insertion sites carrying alleles with different *in vitro* activity (Figure 5B and File S3) showed that potentially inactivating mutations exist in LINE-1 with lower *in vitro* activity than their allelic counterparts. This also implies that intact and *in vitro* active allelic forms of LINE-1s may exist for older insertions - several ‘hot’ LINE-1s in CHM1 belong to the non-Ta subfamily, the oldest of the human-specific LINE-1s. Recent structural variation analyses based on haploid human genome assemblies support this finding that a small set of pre-Ta representatives possibly remain active in the human genome (50). Based on the different *in vitro* activity of LINE-1s alleles at three loci, Seleme *et al*. proposed a model in which LINE-1s accumulate inactivating mutations over time [26]. Our data agrees with this model with additional support of all such cases in a whole haploid genome. We conclude that the collection of *in vitro* active LINE-1s is variable among humans, and some of the active LINE-1s are ‘hidden’ behind the seemingly inactive or even present, not intact allelic forms. From the perspective of LINE-1 evolution, lineages remain active in the human population for much longer than we thought before, with the persistence of active, low-frequency alleles of old LINE-1s.

Previously, *in vitro* activity of LINE-1s has been widely used as a surrogate of their retrotransposition rate in the genome. Although this surrogate fits the context of studying the biology of LINE-1s, the subtle difference between *in vitro* activity and *in vivo* fitness underlies key information of the adaptation of recently inserted LINE-1s. Potential sources of difference between *in vitro* activity and *in vivo* fitness include but are not limited to epigenetic regulation, variable expression of restriction factor, and human population stratification. Previous approaches have used clustering and consensus sequences to estimate *in vivo* fitness based on copy numbers [1,50,51]. However, these studies mostly focused on long timescale changes in TE sequence and abundance using purely computational methods.

We found that the correlation between LINE-1 *in vitro* activity and *in vivo* fitness is dependent on the frequency of the LINE-1 in the population: significant correlation only exists in the polymorphic LINE-1s (Figure 5A). Furthermore, outliers with discrepant *in vitro* activity and *in vivo* fitness are observed within the polymorphic LINE-1s. Comparison of the sequence and *in vitro* activity of LINE-1s at the same allele suggests that most of *in vivo* fit but *in vitro* inactive LINE-1s resulted from assaying the inactive (likely rare) allelic form (Figure 5B). Phylogenetic analysis of all *in vivo* fit and additional *in vivo* unfit but *in vitro* active LINE-1s shows that *in vivo* fit LINE-1s and form distinct supported clusters (Figure 5C, magenta dots), hinting the potential key mutations that are evasive to host regulation and gave rise to the *in vivo* fit LINE-1 lineage. Moreover, multiple supported phylogenetic clusters are found within the *in vivo* fit group, suggesting the diversification of these LINE-1s, and offering the currently *in vivo* fit LINE-1s ability to further adapt to the genome environment. Besides, 14/34 of the *in vivo* fit LINE-1s in CHM1 are either absent or have a non-intact counterpart in CHM13, and have an average frequency of 41.4%, suggesting that the adapting LINE-1s are sparsely distributed in the human population. Together, we conclude that each person carries a different set of active and adapted LINE-1s.

Our study offers a nearly complete investigation of all LINE-1s in a haploid genome. Such investigation allowed us to capture all young LINE-1s in an ‘actual’ genome, rather than the inference based on the ‘average’ reference genome. Based on the LINE-1 sequences in two haploid genomes, we found that LINE-1 allelic variation is much higher than previously estimated and inferred that each person has a unique set of intact or active LINE-1s. Our data is a pilot to much finer scale analyses to predict and measure the activity load of LINE-1s in diverse human genomes. We found that the correlation between the *in vitro* activity and *in vivo* fitness of LINE-1s is dependent on their frequency in the population, defining the conditions required to impute the properties of LINE-1s that were only previously available through wet lab experiments. The rare outliers with ‘conflicting’ *in vitro* activity and *in vivo* fitness revealed the persistence of older LINE-1 lineages and emerging adaptations of the youngest LINE-1s to the genomic and cellular context of the host. Because of LINE-1s’ capability to reorganize the host genome, our study of young LINE-1s at such refined resolution is a significant advancement in understanding human genetic variation.

Although the total number of LINE-1s may appear similar between closely related genomes, the collection of intact (potentially ‘active’ or ‘functional’) LINE-1s can be dramatically different. Given that LINE-1s, particularly the young and polymorphic ones, are known to be either causal to or associated with a slew of human diseases [6], such variation in LINE-1s could be a major factor to consider when inferring the ‘risk score’ of individuals for these diseases.

## Materials and Methods

### Retrieving LINE-1 from the haploid and reference genomes

LINE-1s were identified according to the RepeatMasker annotation from CHM1 and CHM13 assemblies (GenBank assembly accession: GCA_001297185.2 and GCA_000983455.2). BLAST searches of CHM1 and CHM13 used L1.3 (GenBank accession number: L19088) as a query. The RepeatMasker annotation was filtered for keyword ‘L1’ in the ‘matching repeat’ column and converted to bed format. Subsequently, LINE-1 sequences were retrieved from the genome according to the bed file using the ‘subseq’ function of seqtk (https://github.com/lh3/seqtk). GRCh37 and GRCh38 reference genome sequences and RepeatMasker annotations were downloaded from the annotations of GenBank assembly GCA_001297185.2 and GCA_000983455.2 and processed in a similar manner to the CHM genomes to find LINE-1s. LINE-1s on the ALT contigs of the reference genomes were manually inspected for their corresponding chromosomal location if possible and assigned as an alternative LINE-1 allele of that chromosomal location in the reference genome.

### Intact LINE-1 identification

Intact LINE-1s were identified following our previous protocol [52]. Full-length LINE-1s (Files S1 and S2) were found by filtering the RepeatMasker annotation of the CHM genomes requiring the length of the annotated LINE-1 sequence to be equal to or longer than 5,000 bp. LINE-1 ORFs were found by using EMBOSS [53] ‘getorf’ function on full-length LINE-1s with ‘ -find 1’ setting to return the translated sequences of the ORFs. The translated ORFs were subsequently searched using BLASTp with the translated ORFs of L1_RP_ (GenBank accession number: AF148856). For each of the full-length CHM1 and CHM13 LINE-1, a custom perl script processed the BLASTp output to find the ORF of the LINE-1 that forms the longest alignment to the ORF1 and ORF2 protein of L1_RP_. LINE-1s with intact ORFs were identified in the distribution of the longest called ORFs of each LINE-1 that align to the reference ORFs, which correspond to ORF1 length of 338 codons and ORF2 length of 1,275 codons. Singletons near these ORF lengths were manually inspected to find additional ORFs that align to the full length of the L1_RP_ reference ORFs.

### Sequencing of LINE-1s with potential sequencing errors

LINE-1s containing frame-shifting mutations were identified by aligning all annotated LINE-1s in CHM genomes to the protein sequences of L1_RP_ ORFs using the setting of ‘-F15’ of LAST (http://last.cbrc.jp). Number of frame-shift mutations were counted for each LINE-1 using a perl script, and LINE-1s with one or two frame-shift mutations were kept for further analyses. Status of each of these LINE-1 were then compared to the status of the LINE-1 in the reference genome (GRCh38) and the other CHM genome. Under the assumption that sequencing errors are unlikely to happen at the same site of different genomes, we focused on re-sequencing the LINE-1 that contain frame-shifting mutations that are not shared with the reference genome or the other CHM genome. LINE-1s were PCR amplified from selected CHM1 BACs that contain the target LINE-1 and only a single LINE-1 (BPRC, https://bacpacresources.org). PCR products were purified and region containing the frame-shift mutilations were sequenced at GENEWIZ. Using the same sequencing strategy, we also sequenced the top five CHM1 LINE-1s in terms of allelic difference to CHM13.

### Synteny of LINE-1 in haploid and reference genomes

CHM1 and CHM13 genome sequences were aligned, chromosome by chromosome, to the GRCh37 and GRCh38 reference genomes using lastz [54] under the setting of ‘--notransition --step=20 --format=lav’. The alignments were then processed using lavToPsl, axtChain, and chainMergeSort functions of the UCSC genome browser utilities (http://hgdownload.soe.ucsc.edu/admin/exe/) with default settings. LINE-1 coordinates on CHM1 and CHM13 were subsequently converted to GRCh37 and GRCh38 coordinates with the default setting of the liftOver tool of the UCSC genome browser utilities and the processed chain file mentioned above. For the LINE-1s that could not be directly lifted-over with the default liftOver setting, we took the sequences that are 2,000 bp from each end of LINE-1s and lifted them over to the reference genomes. For this step, the lifted-over coordinates of both extended ends were in the same neighborhood of the target reference genome for almost all LINE-1s. We were unable to obtain coordinates for a small subset of LINE-1s flanked by repetitive sequence using this method, and so are unable to assign a genomic position and to pair possible allelic variants; one exceptional LINE-1 flanked by a repeat-rich region had enough unique sequence on both sides to assign as allelic in the CHM1 and CHM13 genomes but still could not be assigned to a genomic coordinate (Table S2, row 183).

### Population frequencies of intact LINE-1s

To retrieve the frequency of intact CHM1 and CHM13 LINE-1s in the general human population. We utilized data from the thousand genome project (KGP) [37,55], euL1db [46] and two complementary studies [22,47]. CHM1 and CHM13 LINE-1s were lifted to hg19 coordinates using the UCSC liftover tool. Structural variation calls including the KGP phase 1 [55], KGP phase 3 [37] and Wong *et al*. [47] were downloaded from http://dgv.tcag.ca/dgv/docs/GRCh37_hg19_supportingvariants_2016-05-15.txt. Where a LINE-1 is present in the reference genome, the deletion calls that overlapped with the intact CHM LINE-1s were found using the ‘ -f 0.9 -r’ setting of the ‘intersect’ function of bedtools (10.1093/bioinformatics/btq033). Where a LINE-1 is absent in the reference genome, the overlapping insertion calls in KGP were found using the default setting of the ‘intersect’ function of bedtools. Frequency of euLINE-1db (10.1093/nar/gku1043) LINE-1s were calculated by the ratio of the number of individuals to the total individuals included in the corresponding study for each ‘mrip’ insertion. Overlapping LINE-1s between CHMs and euLINE-1db were then identified using the default setting of the ‘closest’ function of bedtools. Because the data shows that most of the non-reference LINE-1s overlapped with mrip entries of euLINE-1db, only LINE-1s with 0 distance between CHM and euLINE-1db were considered the same LINE-1. LINE-1 insertion calls of Iskow *et al*. [22] were also intersected to the intact CHM LINE-1s using bedtools. A perl script was then applied to filter out the singleton calls that correspond to only one individual in the population. Frequency of LINE-1s were then calculated based on the ratio of individuals that carry the corresponding insertion or deletion to the total number of individuals included in the study. For each LINE-1, when frequency data is available from multiple resources, the resource with a larger population was always used in the later analyses; when a new insertion was not identified in any of the abovementioned population level studies, LINE-1s were inferred to have < 0.1% frequency in the population; when a CHM LINE-1 overlaps with a reference LINE-1 but no deletions in the population, it was inferred to be fixed or have 100% frequency in the population.

### Cloning of intact LINE-1

We isolated DNA from the BACs we had identified as containing the intact LINE-1s, then digested it using the restriction enzyme SalI (NEB). After digestion we amplified the LINE-1s using a set of primers that allowed us to obtain as much as possible of the full LINE-1 element from each BAC and appended linkers to each end of the products to allow for Gibson assembly into the AscI site of pYX-New-MCS which is a modified version of pYX017 with a BamHI-AscI-BstZ17I cloning site between the existing FseI and PciI sites. This vector allows excision of a LINE-1 using AscI. PCRs were performed using NEB Q5 polymerase with each 50uL volume reaction containing 10uL of 5X Q5 reaction buffer, 1uL of 10mM dNTPs, 2.5uL of 10uM forward and reverse primers, 1uL of template DNA (~100ng/uL), 0.25uL of Q5 polymerase, and 32.5uL water. The thermo cycler protocol is: 1) 98°C for 30 seconds (s), 2) 98°C for 10s, 3) 49°C for 30s, 4) 72°C for 360s, 5) repeat steps 2-4 29 additional times, 6) 72°C for 600s, 7) hold at 4°C. We gel purified all successful PCR products using the Zymoclean Gel Purification Kit (Zymo Research), then quantified purified PCR products using a Nanodrop and inserted them into AscI-digested pYX-New-MCS using Gibson assembly via the NEB HiFi DNA Assembly kit. In order to identify assembled vectors that contained the intact LINE-1s we had inserted, we first inserted the products of the Gibson assemblies into *E. coli* (NEB), grew colonies overnight on LB Agar with ampicillin, and isolated 10 colonies per LINE-1. After growing the selected colonies overnight, we isolated the vector, then performed PCR using LINE-1 specific primers to confirm presence of the correct insert within the vector.

### In vitro assay for LINE-1 activity

For each LINE-1 we selected a total of 3 vectors that had been PCR confirmed to have the correct insert to use for our activity assay. On day 1 of the assay, we transfected approximately 200ng vector into approximately 25,000 293T cells per well in white 96 well tissue culture plates (Genesee Sci) using the TransIT-LT1 transfection reagent (Mirus Bio) and placed the plates in the incubator overnight. The following day, day 2, we spun down the plates to seat the cells, removed the transfection media, and added 250 uL per well of DMEM with 2.5 ug/uL puromycin. We then returned the plates to the incubator until day 5, when we removed them from the incubator, spun them to seat the cells, and removed the media. After removal of the media, we added 25 uL Dulbecco’s PBS and 25 uL of reagent 1 from the Dual-Glo luciferase kit (Promega Inc.) and mixed well to lyse the cells. After approximately 10 minutes we measured *Firefly* luciferase activity with a plate reader. Then we added reagent 2 from the Dual-Glo kit, waited another 10 minutes, and measured *Renilla* activity.

Each 96 well plate contained 4 wells of positive control (L1_RP_ in pYX017) and 4 wells of negative control (pYX-New-MCS, empty) for standardization and to assess the quality of each plate. Additionally, each plate contained 8 wells each of 11 vectors. We randomized which LINE-1s were present on each plate and the order in which they appeared on the plate, but all wells for any given vector containing a LINE-1 were in the same column on a plate. Each vector containing a LINE-1 was assayed at least 3 times, on at least 2 different plates, and spread out over multiple days to minimize the influence of day and plate effects. In total, each vector containing a LINE-1 was assayed in at least 24 wells, and each LINE-1 was assayed in at least 72 wells (3 vectors containing that LINE-1, each assayed in at least 24 wells). The average luciferase activity reading of the positive and negative controls were taken as 100% and 0% LINE-1 activity reference. Luciferase measurements for each LINE-1 were converted to percent activity based on these references.

### Collection of LINE-1 alleles

LINE-1 alleles were collected by mining the long read sequencing data of the GIAB project [45]. We used the data of the reference individual (HG001, also known as NA12878), the Ashkenazim trio (HG002, HG003 and HG004) and a Chinese individual (HG005). PacBio reads from each library were aligned to the L1_RP_ reference using BLAST. Any read that hit L1_RP_ were further mapped to each LINE-1 allele of interest (LINE-1 sequence at the locus plus flanking 2kb). Reads spanning genome-LINE-1 junctions were extracted and aligned for each individual and each LINE-1 allele. Because each individual can have up to two LINE-1 alleles at a given locus, reads at each locus were manually sorted into separate alleles based on their shared changes. Consensus sequence of reads belonging to each LINE-1 allele were taken to represent the allele. LINE-1 alleles were validated in the trio data based on the offspring acquiring an allele from each parent.

## Acknowledgements

This work was supported by a grant from the National Institutes of Health (R35GM142733-01 to RNM and T32GM007270 to RPD). We thank Miriam Rosenberg, John Huddleston, Holly Wichman, Michael Metzger, members of the Metzger lab, and Janet Young for technical advice and critical comments on this manuscript.

