## Supplemental Figures and Legends for "Evolutionary insights from profiling LINE-1 activity at allelic resolution in a single human genome"

**This PDF file includes:**

Figures S1 to S4
Legends for Tables S1 and S2
Legends for File S1 to S4

**Other supporting materials for this manuscript include the following:**

Table S1 and S2
File S1 to S4

Figure S1

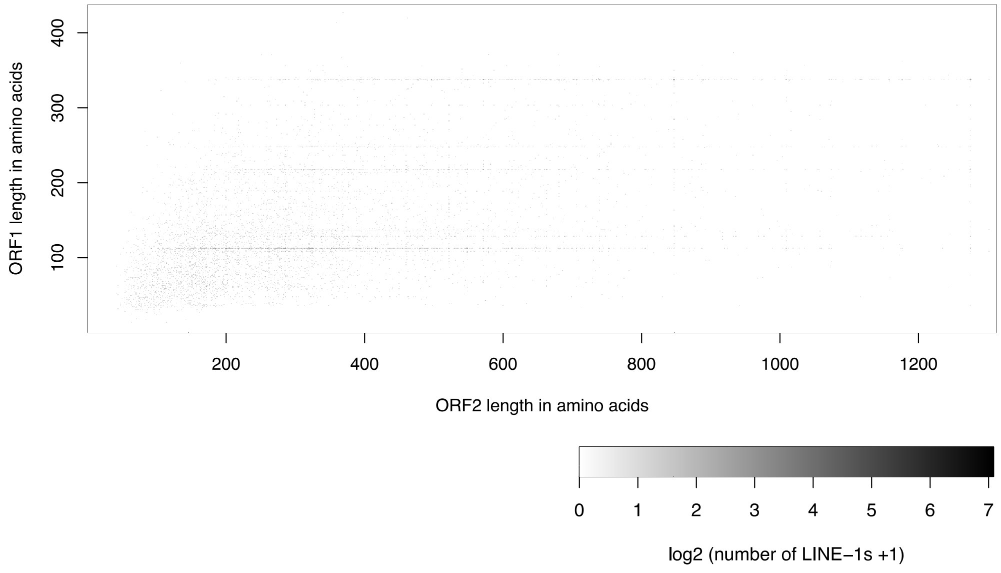


**Figure S1. Heatmap of translated ORF lengths in full-length LINE-1s from CHM1.** Matrix of all called translated ORF lengths in the complete set of full-length LINE-1 sequences (>5000bp). Each pixel represents the number of full-length LINE-1s with the corresponding length of ORF1 (vertical axis) and ORF2 (horizontal axis). Many full-length sequences have either an intact ORF1 or ORF2, but only 148 have both intact ORFs that align along the entire length of a reference amino acid sequence ($L1_{RP}$), mostly concentrated in the pixel in the upper-rightmost region of the heatmap.

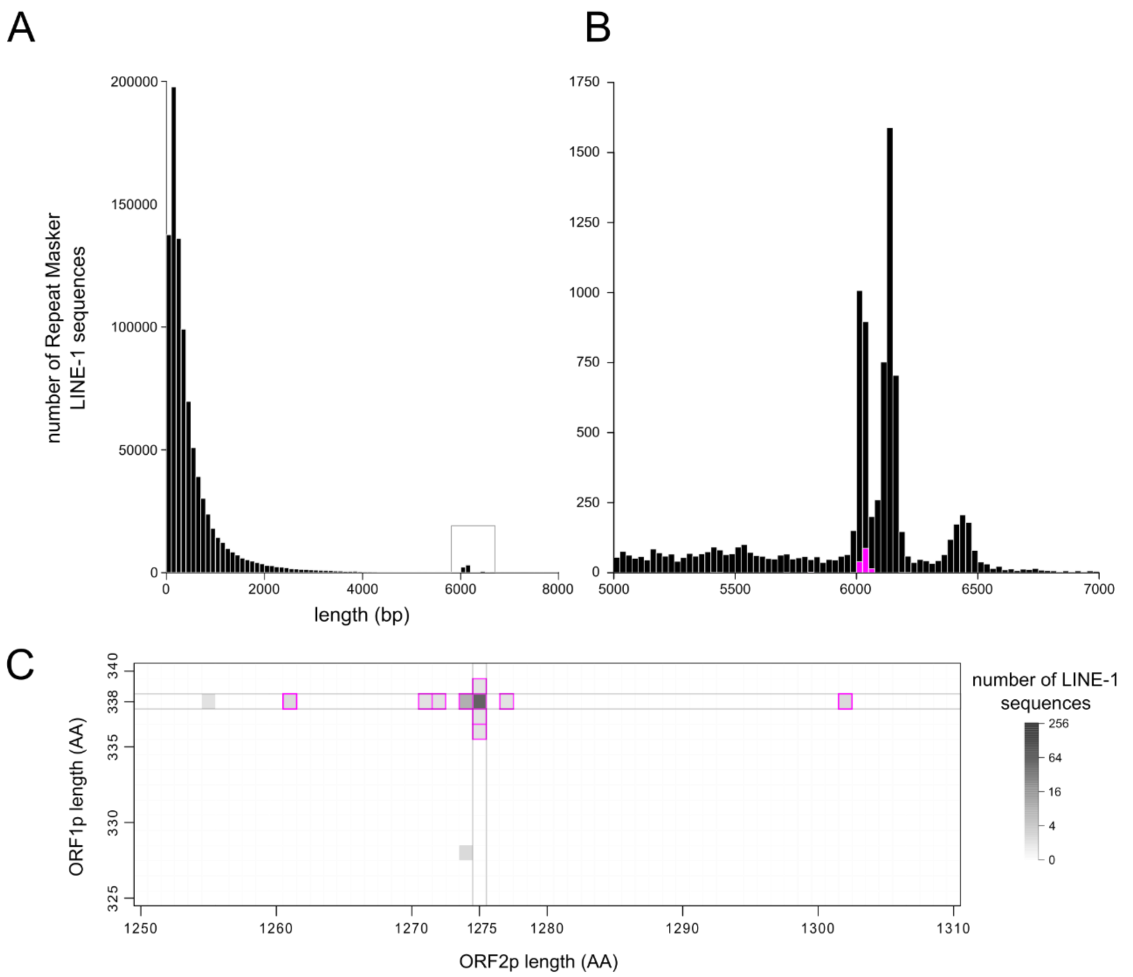


**Figure S2. Identification of intact LINE-1s in CHM1.** A. A histogram of the lengths of sequences in CHM1 that were called by Repeat Masker as LINE-1 shows most sequences are less than 1,000 bp, consistent with the tendency of LINE-1s to be 5' truncated. B. A zoomed-in view of the boxed region containing the peak around 6,000 bp in panel A. This region contains the full-length LINE-1 copies within two sub-peaks: a larger one at around 6,100 bp and a smaller peak at around 6,400 bp. The sequences within these two peaks are distinguished by the presence or absence of a 129bp sequence in their 5'UTR which contains a binding site for ZNF93, a transcriptional repressor of LINE-1s. Intact LINE-1 sequences are highlighted in magenta and found exclusively in the 6,100 bp peak. C. A zoomed in subset of Figure S1 showing the number of sequences with the indicated translated ORF1 and ORF2 lengths showing the pixels containing intact LINE-1 sequences. Within the full-length set of LINE-1 sequences, only ~150 sequences (magenta-outlined pixels) encode putative ORFs that align along the entire length of the ORF1p and ORF2p sequences of a reference element ($L1_{RP}$).

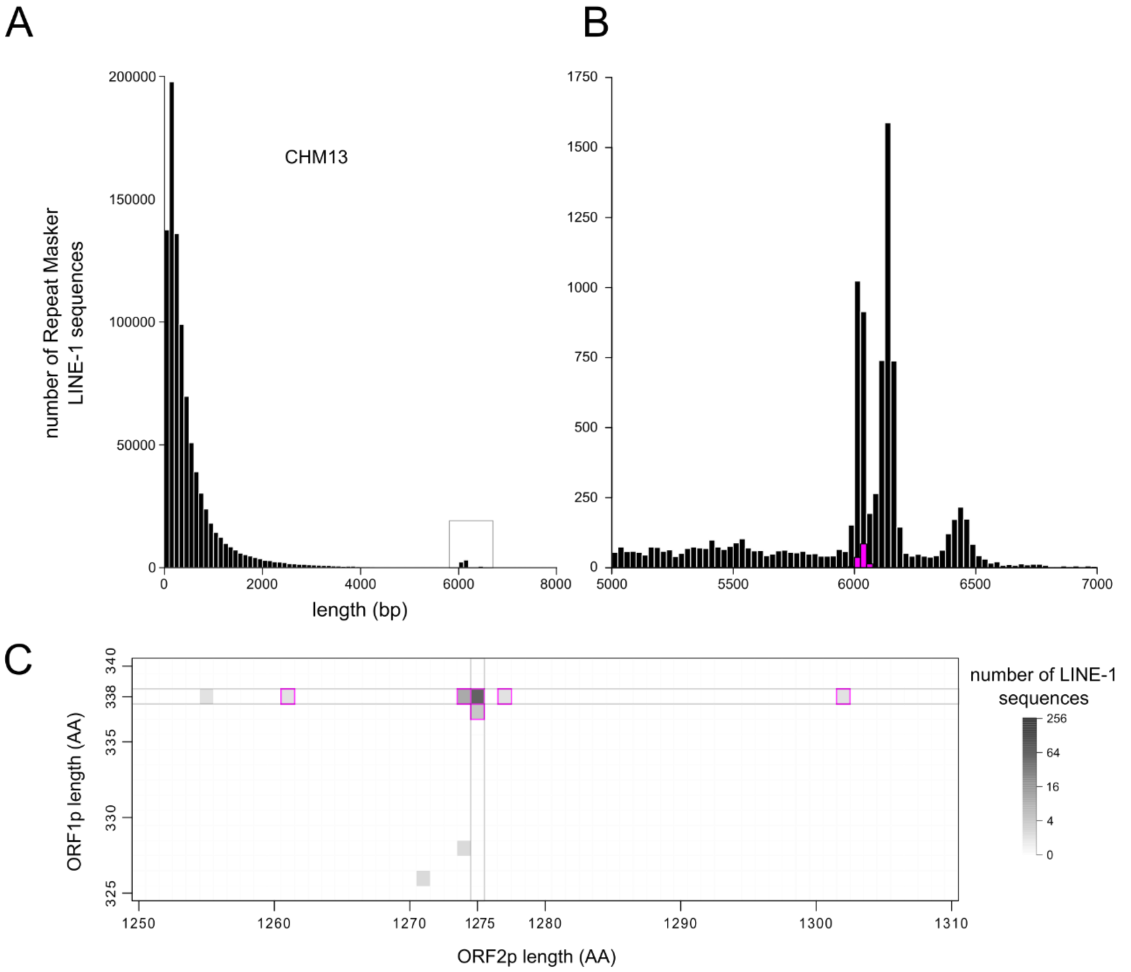


**Figure S3. Identification of intact LINE-1s in CHM13.** A. A histogram of the lengths of sequences in CHM13 that were called by Repeat Masker as LINE-1 shows most sequences are less than 1,000 bp, consistent with the tendency of LINE-1s to be 5' truncated. B. A zoomed-in view of the boxed region containing the peak around 6,000 bp in panel A. This region contains the full-length LINE-1 copies within two sub-peaks: a larger one at around 6,100 bp and a smaller peak at around 6,400 bp. The sequences within these two peaks are distinguished by the presence or absence of a 129bp sequence in their 5'UTR which contains a binding site for ZNF93, a transcriptional repressor of LINE-1s. Intact LINE-1 sequences are highlighted in magenta and found exclusively in the 6,100 bp peak. C. A zoomed in subset of Figure S1 showing the number of sequences with the indicated translated ORF1 and ORF2 lengths showing the pixels containing intact LINE-1 sequences. Within the full-length set of LINE-1 sequences, only ~150 sequences (magenta-outlined pixels) encode putative ORFs that align along the entire length of the ORF1p and ORF2p sequences of a reference element ($L1_{RP}$).

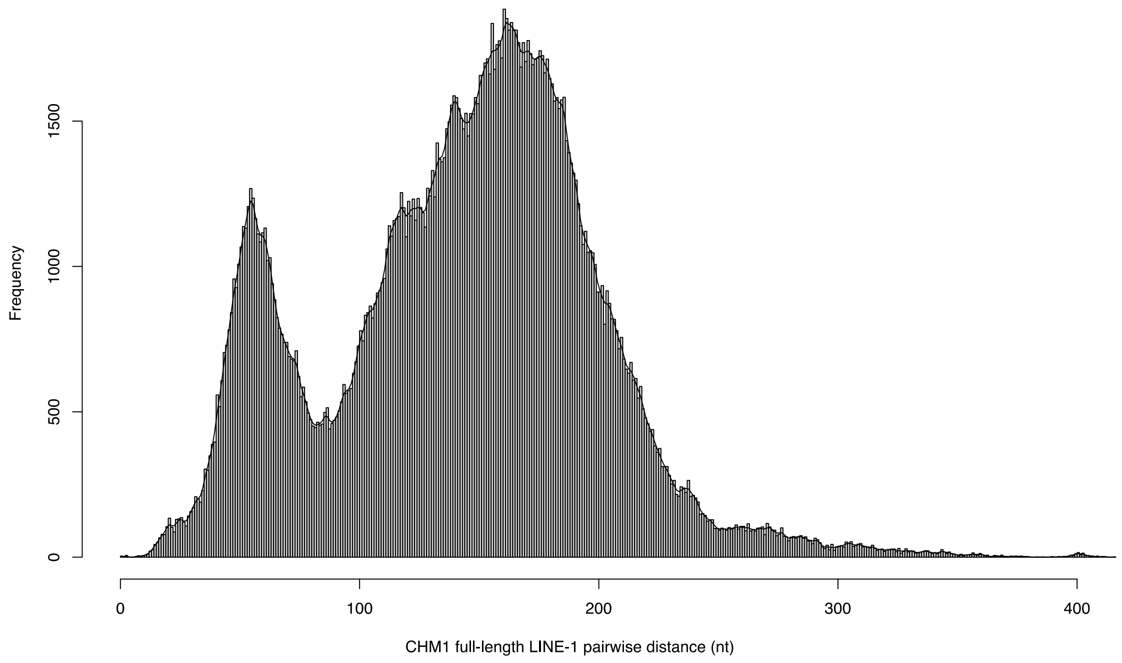


**Figure S4. Combined distribution of pairwise distances of full-length LINE-1 sequences in CHM1.** The curve is generated based on kernel smoothing (bandwidth=3 in using the ‘ksmooth’ function of R) of the distribution. Two major gaps at 28 and 82 substitutions were observed on the curve and used as the cutoff to separate the *in vivo* fit (<28), unfit (28-82), and very old LINE-1 (>82) categories.

**Table S1. A comparison of the intact LINE-1s between GRCh38, CHM1, and CHM13 genome assemblies.** Counts of the number of LINE-1s that are intact, present but not intact, and absent in one genome assembly compared to the same categories in a second genome assembly are shown. NA represents a category where there are no intact LINE-1s in the compared categories of the two genome assemblies.

**Table S2. Analysis of intact LINE-1 alleles from GRCh38 and CHM genome assemblies.** Coordinates of LINE-1s that are intact in either CHM1, CHM13, or GRCh38 (hg38) are shown. GRCh37 (hg19) coordinates are based on liftOver from GRCh38 coordinates. 'Intact' represents the LINE-1s with intact ORFs, 'short' represents the LINE-1s that are present at the insertion site but the ORFs are no longer intact, and '.' indicates that the site does not contain the LINE-1 insertion. LINE-1 subfamilies are classified based on characteristic sites reported by Boissinot *et al.* 2000 (ref 49 of main text). Approaches used to calculate LINE-1 activity (*in vitro*), population frequency, and number of near neighbors (*in vivo* fitness) are described in the Materials and Methods section of the main text.

**File S1. Nucleotide sequences of CHM1 intact LINE-1s.** Sequences are named based on their coordinate in the CHM1 assembly.

**File S2. Nucleotide sequences of CHM13 intact LINE-1s.** Sequences are named based on their coordinate in the CHM13 assembly.

**File S3. Codon-based alignment of LINE-1 nucleotide sequences to identify potentially inactivating mutations in LINE-1 alleles.** The first ten sequences (L1RP-L1PA8A) are the consensus ancient LINE-1 sequences from Khan *et al*. 2006. The following sequences are LINE-1 alleles, separated by empty sequences labeled to denote the *in vitro* activity difference between alleles. Sequence names starting with ‘LJII’ are from CHM1; sequence names starting with ‘LBHZ’ are from CHM13; sequences named with chromosome and coordinate are from GRCh38; sequences named with GenBank accession numbers are from the nt or fosmid database of NCBI; sequences with ‘A’ at the beginning of their names are the consensus alleles from GIAB long reads corresponding to LINE-1 insertion sites. The last sequences annotated ‘other active L1s’ are other *in vitro* active LINE-1s measured in either this study, Brouha *et al*. 2003, or Beck *et al*. 2010.

**File S4. Nucleotide alignment of LINE-1 sequences to identify cluster-defining mutations.** Sequence names indicate their status/lineage and position in the genome. Separated by '_', the first field of the sequence name indicates status of the sequence: 'ActiveUnfit' means that the LINE-1 is *in vitro* active but *in vivo* unfit; sequences not named 'ActiveUnfit' are *in vivo* fit, with the text in this field indicating the LINE-1 subfamily they belong to. The second field contains the last four digits of the scaffold number in CHM1 they belong to. The third field labels the starting coordinate of the LINE-1 on the scaffold.
